# Ketamine anesthesia induces gain enhancement via recurrent excitation in granular input layers of the auditory cortex

**DOI:** 10.1101/810978

**Authors:** Katrina E. Deane, Michael G. K. Brunk, Andrew W. Curran, Marina M. Zempeltzi, Jing Ma, Xiao Lin, Francesca Abela, Sümeyra Aksit, Matthias Deliano, Frank W. Ohl, Max F. K. Happel

**Affiliations:** Leibniz Institute for Neurobiology, D-39118 Magdeburg, Germany; Graduate School of Life Science, Julius Maximilians University, D-97074 Würzburg, Germany; University of Pisa, I-56126 Pisa, Italy; Institute of Biology, Otto von Guericke University, D-39120 Magdeburg, Germany; Center for Behavioral Brain Sciences (CBBS), 39106 Magdeburg, Germany

## Abstract

The N-methyl-D-aspartate (NMDA) receptor antagonist, ketamine, is commonly used as an anesthetic agent and has more recently gained attention as an antidepressant. Ketamine has been linked to increased stimulus-locked excitability, inhibition of interneurons, and modulation of intrinsic neuronal oscillations. However, the functional network mechanisms are still elusive. A better understanding of these anesthetic network effects may improve upon previous interpretations of seminal studies conducted under anesthesia and have widespread relevance for neuroscience with awake and anesthetized subjects as well as in medicine. Here, we investigated the effects of anesthetic doses of ketamine (15mg kg^−1^/h i.p.) on the network activity after pure tone stimulation within the auditory cortex of male Mongolian gerbils (*Meriones unguiculatus*). We used laminar current source density (CSD) analysis and subsequent layer-specific continuous wavelet analysis to investigate spatiotemporal response dynamics on cortical columnar processing in awake and ketamine-anesthetized animals. We found thalamocortical input processing within granular layers III/IV to be significantly increased under ketamine. This effect on early thalamocortical input processing was not due to changes in cross-trial phase coherence. Rather, the layer-dependent gain enhancement under ketamine was attributed to a broadband increase in amplitude reflecting an increase in recurrent excitation. The time-frequency analysis is further indicative of a prolonged period of stimulus-induced excitation possibly due to a reduced coupling of excitation and inhibition in granular input circuits—in line with the common hypothesis of cortical disinhibition via NMDA-mediated suppression of GABAergic interneurons.

**Statement of significance:** Ketamine is a common anesthetic agent and is known to alter excitability and neuronal synchronicity in the cortex. We reveal here that anesthetic doses of ketamine increase recurrent excitation of thalamic input in the granular layers of the auditory cortex of Mongolian gerbils. This leads to a layer-specific gain enhancement of the time-locked response to external stimuli. Analysis of tone-evoked amplitudes and cross-trial variability of cortical current sources and sinks indicate a mechanism of cortical disinhibition via NMDA-mediated suppression of GABAergic interneurons. Our findings might help to understand the functional mechanisms of the clinical effects of ketamine promoting the development of new therapeutic agents with lower side effects.

## Introduction

In research of cortical function, the use of general anesthesia aims to prevent acute pain and reduce signal noise, while still leaving the sensory information pathway to the neocortex as intact as possible. Gold-standard studies in this field have therefore, historically, been performed under anesthesia or partial anesthesia (e.g. Deweese & Zador, 2003; Heil, 1997b, 1997a; Hubel & Wiesel, 1959, 1962, 1965, 1969; Petersen et al., 2003).While technical advances have increasingly allowed us to explore the physiology of cortical functions in awake and behaving animals, still little is known about their direct comparison, particularly with respect to the interaction between external stimuli and intrinsic cortical dynamics (Pachitariu et al., 2015). A better understanding would help to more accurately interpret findings from both states and therefore have widespread relevance for neuroscience and medicine.

One commonly used surgical anesthetic in systems physiology is ketamine—an N-methyl-D-aspartate (NMDA) receptor antagonist—which enters the open receptor channel inhibiting ionic exchange (Anis et al., 1983; Macdonald et al., 1987). One main effect of ketamine thereby is a persistent increase in cortical glutamate which renders cells more excitable (Miller et al., 2016; Zhang et al., 2019) and disinhibition of the cortex through suppression of GABAergic interneurons (Behrens et al., 2007; Homayoun & Moghaddam, 2007; Schobel et al., 2013). Further, spike sorting is limited under ketamine, potentially due to synchronized population dynamics across large cortical distances (Hildebrandt et al., 2017). Such synchronization has been explored *in vivo* through spectral analysis in various human and animal studies: at anesthetic doses, increased gamma oscillations in cortico-subcortical networks may result from an altered interplay between cortical pyramidal neurons and parvalbumin-expressing interneurons (Grent-’t-Jong et al., 2018; Lazarewicz et al., 2010; Qi et al., 2018; Slovik et al., 2017). Others found differential effects on neuronal oscillations with decreased large-scale beta band activity at both sub-anesthetic (Ma, Skoblenick et al., 2018; Pallavicini et al., 2019) and anesthetic doses (Hertle et al., 2016), and increased theta and decreased alpha EEG (Blain-Moraes et al., 2014; Bojak et al., 2013). One caveat of current studies comparing cortical processing between anesthetic and awake states is that they mainly focus on single or multi-unit level data, or on the macroscopic EEG signal in human research. Studies that characterize the mesoscopic scale—and link the observed effects of ketamine to cortical circuitry processing— are still relatively scarce (c.f. Fitzgerald & Watson, 2019; Michelson & Kozai, 2018).

In this study, we compared tone-evoked current-source density (CSD) distributions in the awake and ketamine-anesthetized (15mg kg^−1^/h i.p.) primary auditory cortex (A1) of Mongolian gerbils (*Meriones unguiculatus*). The feedforward spatiotemporal current flow of tone-evoked activity across cortical layers is highly similar in the awake and ketamine-anesthetized A1, suggesting that canonical cortical population activity is conserved during anesthetic states (Luczak et al., 2012). However, we observed distinct differences in the temporal variability of the overall current flow. A layer-dependent analysis of evoked activity showed that synaptic input in granular layers III/IV was significantly increased under ketamine anesthesia. While signal strength increasee, peak latencies were shorter, and less variable compared to the awake cortex—indicating higher synchrony of tone-evoked cortical inputs. A continuous wavelet analysis revealed a larger time-frequency region of phase coherence across trials to incoming external stimuli mainly in granular layers. However, the layer-dependent gain enhancement of the dominant sensory response early after tone presentation was mainly attributed to a broadband increase in amplitude reflecting an increase in recurrent excitation. Ketamine appears to prolong the period of stimulus-induced excitation due to a reduced coupling of excitation and inhibition in granular input circuits in line with the common hypothesis of cortical disinhibition. Anesthetic doses of ketamine hamper the high synaptic variability accounting for the probabilistic cortical processing mode in the awake brain and might therefore be responsible for its effects on consciousness (c.f. Supp et al., 2011).

## Materials & Methods

All experiments were carried out with adult male Mongolian gerbils (*Meriones unguiculatus*, 4 to 8 months of age, 70-90g bodyweight, total n=20). Experiments were conducted in accordance with ethical animal research standards defined by the German Law and approved by an ethics committee of the State of Saxony-Anhalt.

### Electrophysiological recordings

We recorded activity from the A1 of animals that either received a ketamine/xylazine anesthesia by acute implantation of a recording electrode (n=11) or from awake and passively listening animals with chronically implanted electrodes (n=9).

### Acute recordings and pharmacological silencing of the auditory cortex

The surgical procedure for electrophysiological recording has been previously described in detail (Deliano et al., 2018). Briefly, ketamine/xylazine was administered during surgery and throughout the experiment to maintain a steady anesthesia level. Infusion of 45% v/v ketamine (50 mg/ml, Ratiopharm GmbH), 5% v/v xylazine (Rompun 2%, Bayer Vital GmbH), and 50% v/v of isotonic sodium-chloride solution (154 mmol/1, B. Braun AG) was performed intraperitoneal with an initial dose of 0.004 ml/g bodyweight. After 1-2h of deep anesthesia during the surgery, anesthetic status during the experiment was maintained with infusion at a rate of 15mg/kg^−1^/h.

Anesthetic status was regularly checked (10-15 min) by paw withdrawal-reflex and breathing frequency. Body temperature was kept stable at 34°C. The right auditory cortex was exposed by trepanation and the A1 was located by vascular landmarks. A small hole was drilled on the contralateral hemisphere for implanting a stainless-steel reference wire (Ø 200μm). Animals were head fixed with an aluminum bar, affixed by UV-curing glue (Plurabond ONE-SE and Plurafill flow, Pluradent).

Animals were head-fixed in a Faraday-shielded acoustic soundproof chamber with a speaker located 1 m posteriorly (Tannoy arena satellite KI-8710-32). Local field potentials (LFPs) were recorded with a 32-channel shaft electrode (NeuroNexus A1×32-50-413) implanted in the A1 perpendicular to the cortical surface (c.f. Happel et al., 2010). Recorded LFPs were fed via an Omnetics connector (HST/32V-G2O LN 5V, 20× gain, PlexonInc) into a PBX2 preamplifier (Plexon Inc.) to be pre-amplified 500-fold and band-pass filtered (0.7-300 Hz). Data were then digitized at a sampling frequency of 1000 Hz with the Multichannel Acquisition Processor (Plexon Inc.).

A series of pseudo-randomized pure-tone frequencies covering a range of 7 octaves (tone duration: 200 ms, inter-stimulus-interval (ISI): 800 ms, 50 pseudorandomized repetitions, 65 dB SPL, 7.5 min per measurement, 125 Hz – 32 kHz). We determined the best frequency (BF) as the frequency evoking the strongest response in the averaged granular CSD channels (see below). Stimuli were generated in Matlab (Mathworks, R2006b), converted into analog (sampling frequency 1000 Hz, NI PCI-BNC2110, National Instruments), routed through an attenuator (gPAHGuger, Technologies), and amplified (Thomas Tech Amp75). A microphone and conditioning amplifier were used to calibrate acoustic stimuli (G.R.A.S. 26AM and B&K Nexus 2690-A, Bruel&Kjaer, Germany).

Recordings of tone-evoked responses were taken after recording quality had stabilized: typically 1 h after implantation. After measuring the tonotopic tuning, 20 μl of the GABA_A_ agonist muscimol (8.23 mM muscimol, TOCRIS bioscience, batch no: 9C/107090), dissolved in isotonic sodium-chloride solution, was applied topically onto the cortical surface to silence intracortical contributions of synaptic activity (Ref: Happel et al., 2010). Once the muscimol had diffused through all cortical layers, residual thalamocortical inputs could still be detected by early current sink inputs in layers III/IV and Vb/VIa. Tonotopic tuning of isolated thalamocortical inputs was performed as before cortical silencing.

### Chronic Implantation and in-vivo tonotopy recordings

Chronic implantation of a recording electrode follows similar surgical procedures to the acute implantation. Importantly, the trepanation is kept smaller in order to limit the region of exposed cortex to avoid tissue damage and to achieve stable fixation of the electrode later. A recording electrode with a flexible bundle between shaft and connector (Neuronexus, A1×32-6mm-50-177_H32_21mm) was inserted and an initial recording was conducted in order to guarantee the implantation within A1. Then, the electrode and the connectors (H32-omnetics) both were glued to the skull with a UV-curing glue (Plurabond ONE-SE and Plurafill flow, Pluradent). In order to protect the small exposed region of the cortex, the hole was filled with a small drop of an antiseptic lubricant (K-Y Jelly, Reckitt Benckiser). After the surgical procedure, animals received analgesic treatment with Metacam (i.p. 2mg/kg bw; BoehringerIngelheim GmbH) substituted by 5% glucose solution (0.2ml). Animals were allowed to recover for at least 3 days before the first recording.

Animals were then placed in a single-compartment box in an electrically shielded and sound-proof chamber. Recordings were performed with the head-connector of the animal through a preamplifier (HST/32V-G2O LN 5V, 20× gain, PlexonInc or RHD2132 Omnetics-Intan technologies) and a data acquisition system (Neural Data Acquisition System Recorder Recorder/64, Plexon Inc. or RHD2000 series, Intan Technologies), visualized online (NeuroExplorer, Plexon Inc. and RHD2000 interface GUI Software), and stored. Broadband signals were filtered offline to extract local field potentials (2000 Hz sampling frequency and later down sampled to 1000 Hz). Acoustic stimuli were presented with the same parameters, as in the anaesthetized group. Auditory stimuli were calibrated using a 1⁄2 inch condenser microphone (Brüel&Kjær) and presented with an intensity of 20 dB above a detectable averaged tone-evoked LFP component. Stimuli were digitally synthesized and controlled using Matlab (Mathworks, R2012b) and presented by Presentation® (Neurobehavioral Systems). Stimuli were delivered via an attenuator (g-PAH Guger Technologies), an amplifier (Lehmann Audio) and two electrostatic loudspeakers positioned 5 cm outside both sides of the box.

### Current source density analysis

Based on the recorded laminar local field potentials, the second spatial derivative was calculated in Matlab (Mathworks, R2016a), yielding the CSD distribution as seen in equation 1:

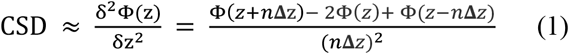

where Φ is the field potential, *z* is the spatial coordinate perpendicular to the cortical laminae, **Δ***z* is the sampling interval, and *n* is the differential grid (Mitzdorf, 1985). LFP profiles were smoothed with a weighted average (Hamming window) of 7 channels which corresponds to a spatial kernel filter of 300 μm (Happel et al., 2010). CSD distributions reflect the local spatiotemporal current flow of positive ions from extracellular to intracellular space evoked by synaptic populations in laminar neuronal structures. Current sinks thereby correspond to the activity of excitatory synaptic populations, while current sources mainly reflect balancing return currents. Early current sinks in the auditory cortex are therefore indicative of thalamic input in granular layers III/IV and infragranular layers Vb/VIa (Happel et al., 2010;Szymanski et al., 2009)

CSD profiles were further transformed by averaging the rectified waveforms of each channel:

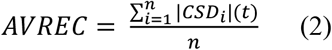

where *n* is the individual channel and *t* is time in ms. This measure gives us the overall temporal local current flow of the columnar activity (Givre et al., 1994; Schroeder et al., 1998).

Based on tone-evoked CSD distributions, we assigned the main sink components to the cortical anatomy as follows: the early dominant sink components are attributed to lemniscal thalamocortical input, which terminates in cortical layers III/IV and Vb/VIa (Figure 1; cf. Happel et al., 2010). Subsequent sink components emerge in supragranular layers I/II, and infragranular layers Va and VIb.

**Figure 1.**
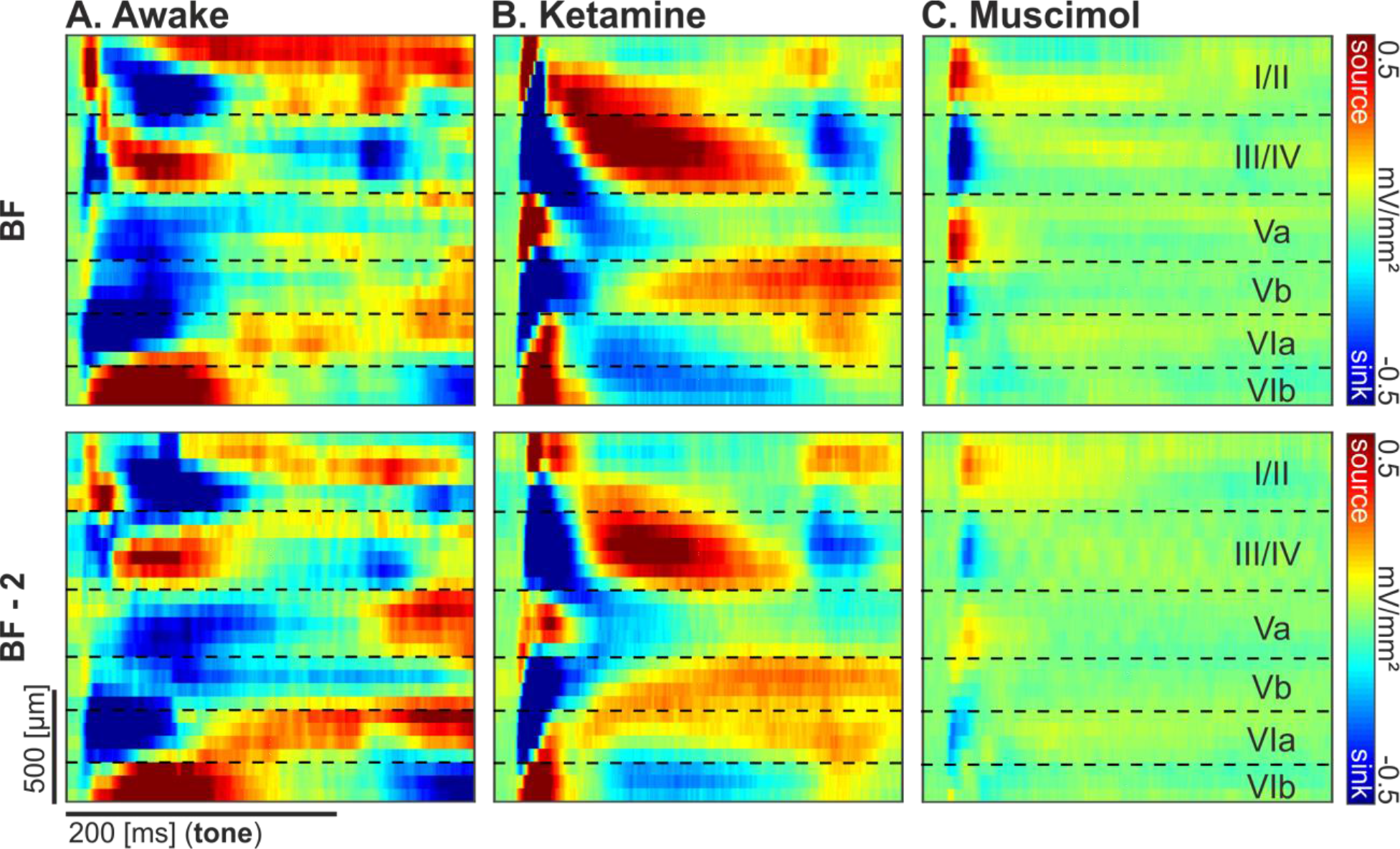
Grand average current-source density profiles. For BF and BF-2 below in ***A***) awake, ***B***) ketamine, and C) muscimol conditions (n=9, n=11 and n=11, respectively). Top, Best frequency response. Bottom, Response 2 octaves below the best frequency. The CSD shows the pattern of temporal processing (ms) within the cortical depth (μm). Representative layer assignment is indicated with horizontal dashed lines corresponding to the layers in C. Pure-tone stimuli were presented for the first 200 ms. Current sinks (blue), represent areas of excitatory synaptic population activity, while current sources reflect balancing currents (cf. Happel et al., 2010).

### Clustermass permutation analysis

To compare the subject averaged CSD for BF stimulation and for frequencies 2 octaves below the BF (BF-2), clustermass permutations were performed between the CSD data of ketamine-anesthetized and awake recordings. At each point of the CSD matrix of observed difference between ketamine – awake subjects, a two-sample student’s *t* test was calculated (viz. 32×600 *t* tests). Each point in the observed difference matrix where *t* was significant of at least p<0.05 was summed and the total value taken as clustermass. The subject measurements were randomized per permutation to compute a clustermass calculation as described above; 1000 permutations were calculated to create a distribution of clustermass values and the observed clustermass total was then compared to this distribution and was found to be significant where the observed total was 1 standard deviation (p<0.05) or more above the mean. This was performed for CSD data in early and late time windows, for selected layers, and for the entire CSD matrix reflecting overall cortical columnar activity. Layers were selected based on CSD profiles overlapping layer rows (i.e. supragranular I/II and thalamocortical input layers III/IV, Vb, and VIa) between the grand averaged anesthetized and awake CSD profiles to avoid analyzing between different cortical layers.

### CSD-derived Tuning curves

Tuning curves of layer-wise tone-evoked activity were centered around the BF response of each respective layer. Tuning curves were calculated for peak latencies and root mean square (RMS) values. Values were detected automatically when the sink activity crossed 1.5 standard deviations below the measured baseline activity. Candidate sink components were detected within each layer for an early time window (1-65 ms after tone onset) and a late time window (66-400 ms after tone onset). Based on the root mean square power, the strongest sink activation was selected per time window.

Tuning curves for average rectified CSD (AVREC) were centered around the BF of the granular thalamocortical sink (layer III/IV). Tuning features at the columnar level were calculated within the first 100 ms of tone presentation as RMS amplitude, peak amplitude, and peak latency.

### Continuous Wavelet Transform

Spectral analysis was performed in Matlab (Mathworks, R2019a) using the wavelet analysis toolbox function CWT for the following variables: animal, condition, stimulus, and recording channel. Important parameters fed into the CWT are as follows: CSD profiles, frequency limits: 5 to 100 Hz (below the Nyquist), and wavelet used: analytic Morse (c.f. Lilly & Olhede, 2012; Olhede & Walden, 2002). For layer-wise wavelet analysis, 3 channels centered around the middle channel of each layer was fed into the CWT and averaged. A trial-averaged scalogram was calculated for each cortical layer and wavelet magnitude—per frequency, per time point—for each subject was computed with equation 3.

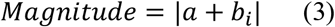

where a + b_i_ represents the complex number output of the trial-averaged CWT analysis. Single trial scalograms were calculated for each animal as well and phase coherence—per frequency, per time point—for each subject was computed with equation 4.

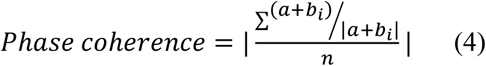

where a + b_i_ represents the complex number output of the single trial CWT analysis (Lachaux, Rodriguez, Martinerie, & Varela, 1999). Magnitude and phase coherence data were averaged pointwise (i.e. frequency and time bins are consistent across averages) for group plots. Clustermass permutations (as above) were performed for the difference between spectral representation in each layer at the BF and two octaves below (BF −2). Frequency bands were split as follows: theta 4-7 Hz, alpha 8-12 Hz, low beta 13-18 Hz, high beta 19-30 Hz, low gamma 31-60 Hz, and high gamma 61-100 Hz. For magnitude calculations: the test statistic for permutation was the student’s *t* test and a Cohen’s D matrix was generated to indicate effect size per frequency at each time point. For phase coherence calculations: the test statistic for permutation was the non-parametric Mann-Whitney U test (toolbox from Matlab file exchange: Cardillo, 2009) and effect size was indicated with the *z* score output as follows:

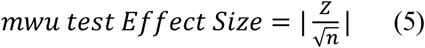

## Results

### Altered spatiotemporal profile in the auditory cortex induced by ketamine anesthesia

We compared the tone-evoked laminar CSD profiles in the primary auditory cortex of Mongolian gerbils under ketamine-xylazine anesthesia, with cortical silencing by topical application of muscimol, and while awake (Figure 1). Comparison of pure tone-evoked grand averaged CSD profiles reveal distinct differences in the spatiotemporal flow of synaptic activity across cortical layers. CSD profiles of both groups (Figure 1A and B) are characteristic of the spatiotemporal flow of sensory information across cortical layers in line with previous findings (Atencio & Schreiner, 2010; Barth & Di, 1990; Sakata & Harris, 2009; Steinschneider et al., 1998; Steinschneider et al., 1992; Szymanski et al., 2009). Approximately 20 ms after pure-tone onset, an initial sink component is detected in granular layers III/IV as well as infragranular layers Vb and Via, all originating from lemniscal feedforward thalamocortical projections to A1 (Happel et al., 2010). From these input circuits, synaptic population activity propagates to supragranular and infragranular layers yielding later sink components (Atencio & Schreiner, 2010; Chen et al., 2007; Sakata & Harris, 2009; Schroeder et al., 1998). This columnar activation is more robust under BF stimulation less when stimulating with a frequency two octaves below (BF-2; cf. Kaur et al., 2005; Happel et al., 2010). Overall, Figure 1 reveals a similar spatiotemporal pattern of auditory-evoked responses with respect to the order of current sinks across layers and time in awake and under ketamine anesthesia. A qualitative comparison reveals a weaker and less expanded early granular sink component and more temporal spread of subsequent current sink components in the awake state. This might be indicative of a ketamine effect on thalamic input activity and early cortico-cortical processing possibly due to less variable and more stimulus time-locked activity.

In order to disentangle thalamic input to the cortex from further cortico-cortical processing, the GABA_A_ agonist muscimol was applied topically to the auditory cortex (Happel et al., 2010). Muscimol silences all intracortically generated activity in the cortex except for the short thalamic input in granular and infragranular layers (Figure 1C). The strength of the early granular sink activity is strongly reduced and shorter under muscimol, reflecting the strong intracortical amplification of early thalamic input under ketamine. Consistently, the spatial width of the granular thalamic sink is reduced under muscimol, indicating that the immediate recurrent excitation in cortical layers III/IV is being blocked by the drug. A comparable reduction of strength and width of tone-evoked activity after silencing intracortical processing can be observed in the early infragranular sink component. Disentangled thalamic input components to the cortical activity pattern are slightly less strong for frequencies two octaves aside of the BF. Qualitative differences between the awake and ketamine group (Figure 2A) are tested for significance via a clustermass permutation test (Figure 2B). Permutation groups are made up of a random selection between awake and ketamine CSD profiles to indicate that the observed differences (Figure 2A) in the two-sample student’s *t* test are above chance level. A comparison of the entire CSD matrix shows that there is significantly more clustermass in the observed measurements than in the distribution of permutations (1000 permutation repetitions). An analysis of cortical layers separately reveals that layer III/IV is highly significantly different across conditions (p<0.001). Less pronounced are significant differences in layers I/II and Vb (p<0.05). The purpose of this permutation test is to validate that effects illuminated by further statistical methods are unlikely to be randomly produced, which is particularly the case for cortical layers III/IV.

**Figure 2.**
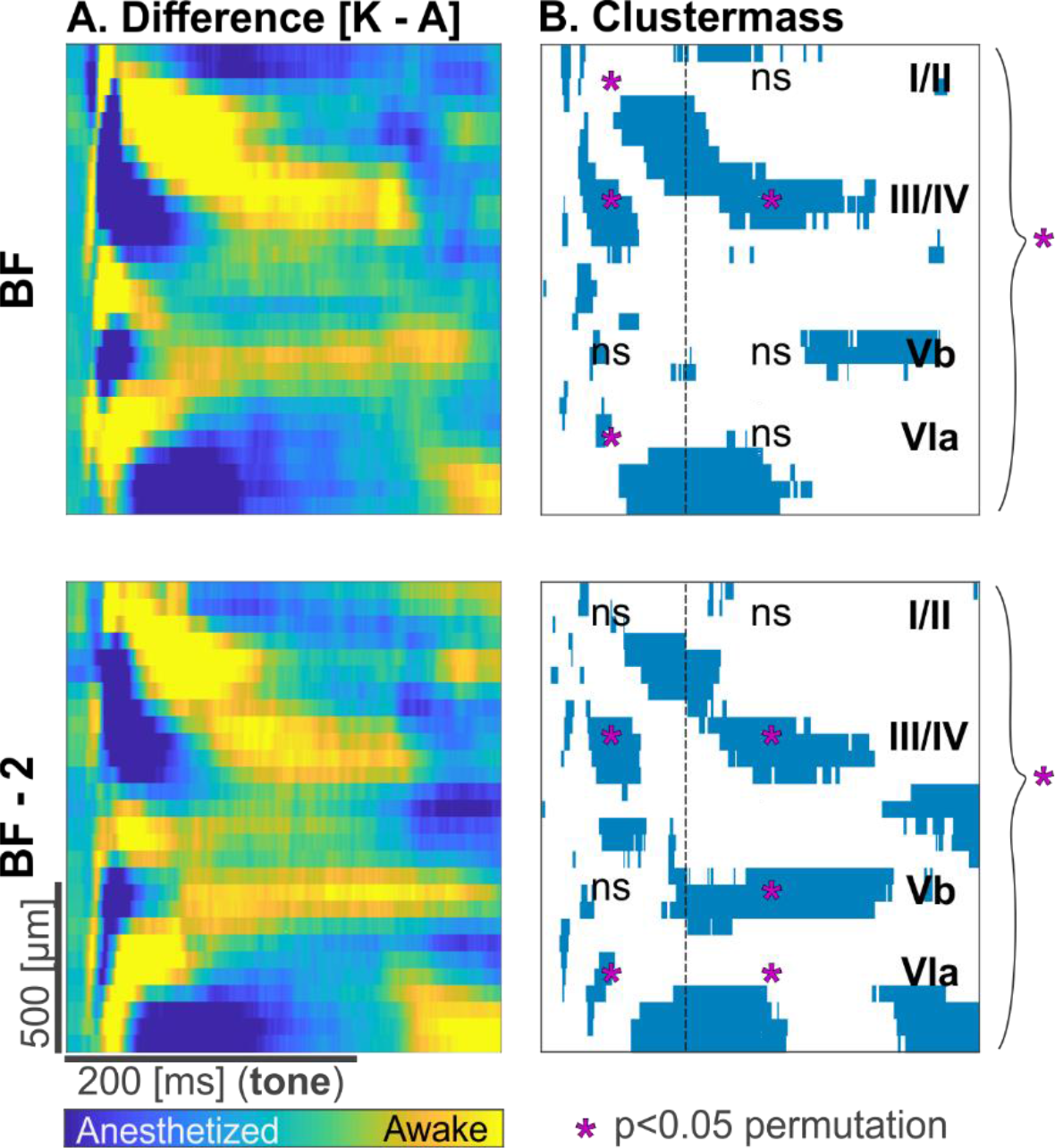
CSD clustermass permutation test. ***A***, The difference (Ketamine - Awake) between the grand averaged CSD profiles for BF and BF – 2 octaves. ***B***, Observed clustermass of significant differences (p<0.05) are plotted in blue (verified by a two-sample student’s *t* test). Significances (p<0.05) of the permutation test are indicated by fuchsia asterisks. Comparisons of the permutation test have been performed for early (1-100 ms) and late (101-400 ms) signals on a layer-wise (I/II, III/IV, Vb, VIa) and columnar level (indicated by braces on the right side).

### Ketamine strengthens stimulus-locked activity by reduced temporal variability of tone-evoked input

The CSD profiles of the awake and ketamine groups indicate temporal differences of tone-evoked activity (Figure 1). We therefore compare the averaged AVREC (Figure 3A) as a measure of the temporal current flow between groups. Ketamine-anesthetized subjects show stronger AVREC onset response peaks. This is indicative of a more stimulus-onset-locked activation of the cortical microcircuitry. Muscimol reduces the response strength but demonstrated a similar pattern of time-locking with BF stimulation. Stimulation with frequencies 2 octaves apart from the BF evoked only a slightly detectable tone-evoked component. Awake subjects show a broadened response after stimulation with both the BF and BF-2. In order to quantify these effects, we further calculated tuning parameters at a single-trial level (Figure 3B). Consistently, under ketamine single-trial peak amplitudes were significantly higher compared to the awake group for all stimulation frequencies (Figure 3B, *left*). The amplitude increase might be due to higher precision of stimulus-locked activity is further implied by the significantly shorter peak latencies in the ketamine group (Figure 3B, *middle*).

**Figure 3.**
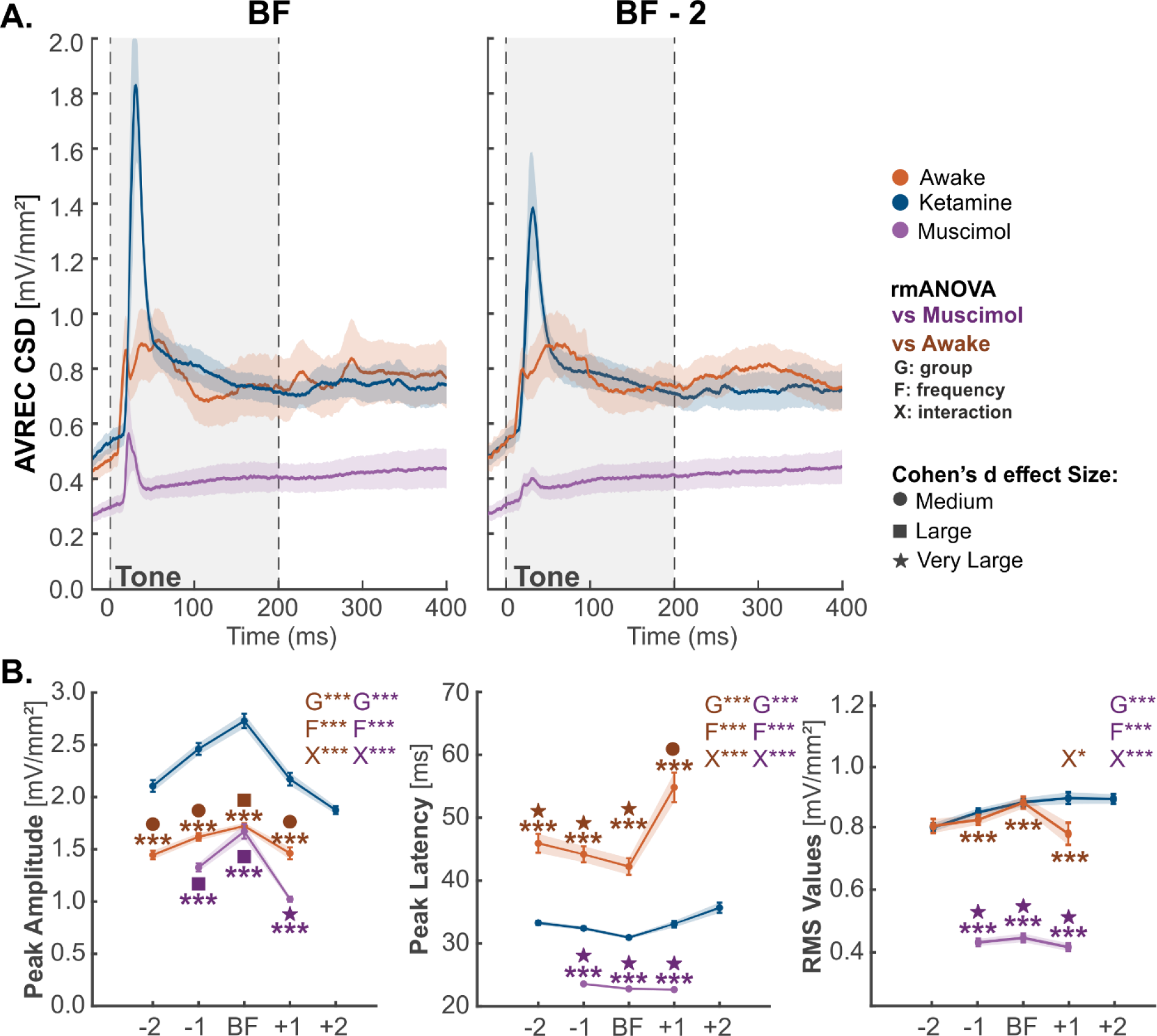
Group-wise comparison between columnar tuning properties. ***A***, A window of 1-100 ms was selected to detect for the most prominent peak within each single trial and the latency and height of that peak was taken for each group (orange: awake, n=9; blue: ketamine, n=11; purple: muscimol, n=11).A: AVREC CSD at BF and BF −2 octaves. ***B***, Tuning curve of peak amplitude (left) and latency (middle) and the RMS of the AVREC trace between 1-100 ms (right). Statistics are based on an rmANOVA, which shows group (G), frequency (F), or interaction (X) effects between both ketamine vs awake (dark orange), or ketamine vs muscimol (dark purple). * =p<0.05. ** =p<0.01, *** =p<0.001. Cohen’s D indicates effect sizes with a circle for medium, a square for large, and a star for very large. A group effect size was also calculated for each stimulus bin.

Due to the stimulus-time-locking under ketamine-anesthesia, muscimol showed even faster peak latencies with less overall strength of the evoked peak amplitude. Muscimol also led to a reduced tuning width around the BF due to a lack of prominent peaks during the detection window of 1-100 ms after tone onset. This can be explained by the silenced intracortical circuitry which normally broadcasts spectral information across cortical space (Kaur et al., 2004). The RMS of the AVREC waveform resembles not the highest activation, but the summed activity over time. Within the first 100 ms after tone onset we observe strong significant difference in the ketamine-anesthetized group before and after muscimol treatment (Figure 3B, *right*). However, the Cohen’s D effect size between the awake and ketamine group shows that the RMS does not differ—indicating that the current flow over time between the awake and ketamine group is rather comparable despite such strong difference between the peak amplitudes and latencies.

The higher overall AVREC peak amplitude, shorter peak latencies, and small Cohen’s D of RMS activity under ketamine anesthesia might be explained by a more synchronized recruitment of synaptic circuits. We therefore assume that the variance of evoked amplitudes would be higher under ketamine anesthesia, while responses peak latencies are more distributed in awake recordings (Figure 4A). To test this, we have applied a Brown-Forsythe test of variance to characterize noticeable differences in variance between timing and amplitude of the peaks of each group. Under ketamine-anesthesia, there is a significantly larger variance in peak amplitude compared to both awake (p<0.001) and with muscimol (p<01; Figure 4B). Conversely, awake subjects have significantly higher variance at peak latencies (p<0.001; Figure 4B). This analysis accounts for the temporally broader shape of the averaged AVREC responses in the awake condition compared to ketamine-anesthetized subjects exhibiting a more stimulus-locked overall activity pattern.

**Figure 4.**
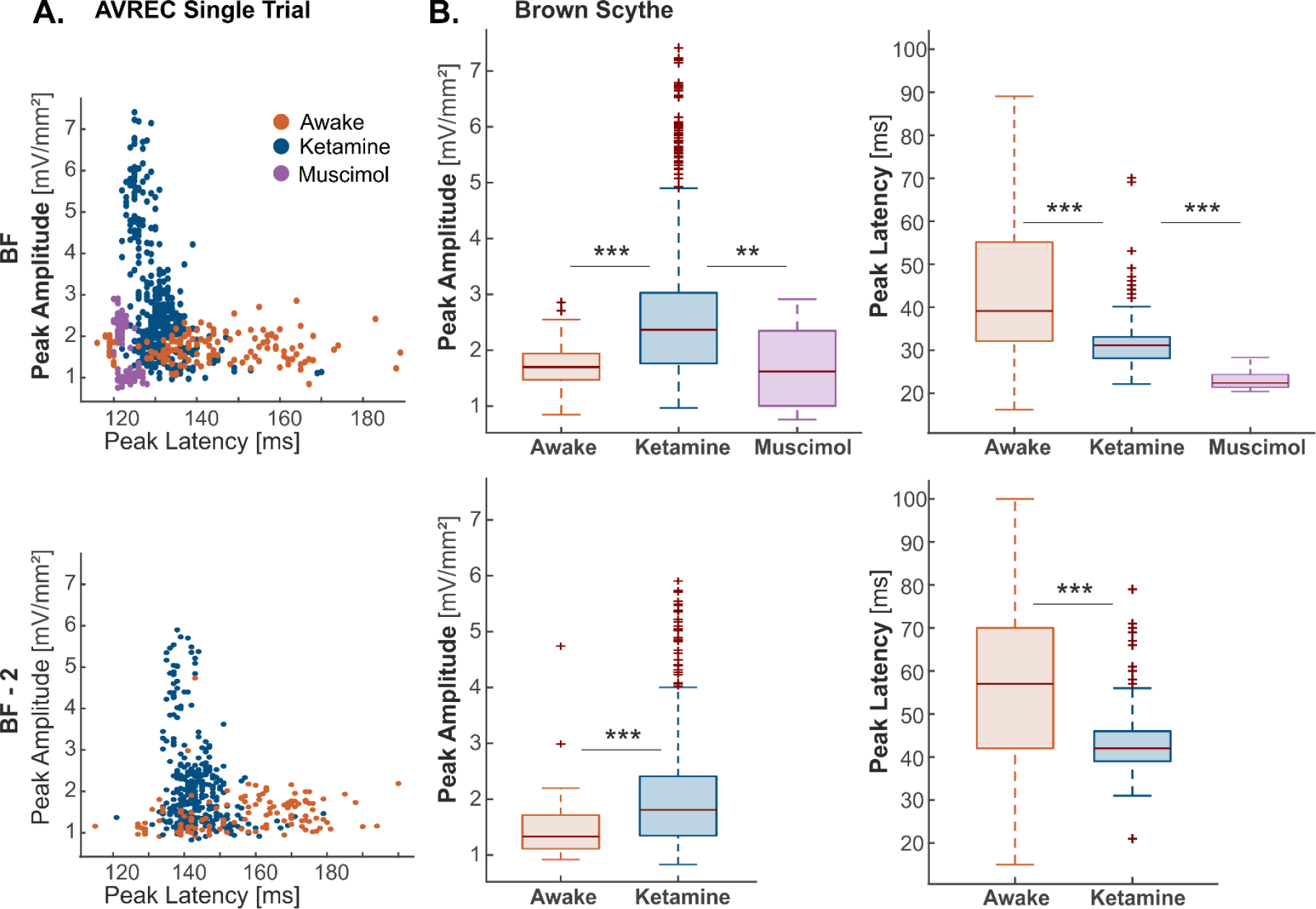
Brown-Forsythe test of variance for peak amplitude and latency. ***A***, Single trial-based scatter plots of amplitude against latency of detected AVREC CSD peaks at BF (top) and BF −2 (bottom). ***B***, Brown-Forsythe variance plotted as boxplots for awake (orange, n=9), ketamine (blue, n=11), and muscimol (purple, n=11) groups for peak amplitudes (left) and latencies (right). Boxes represent the 25%-quartiles, the bar represents the median, and whiskers represent the full range of data except of outliers plotted as crosses. Note that cortical silencing via muscimol reduces measurable activity in non-BF stimulations. *** = p<0.001, ** = p<0.01

### Ketamine-induced gain increase is due to amplitude-effects on granular input layer activity

In order to identify the source of the overall columnar differences, laminar tuning curves were calculated for various parameters of detected current sinks for each animal at thalamocortical input layers III/IV, Vb, and VIa, and supragranular layers I/II (Figure 5). The main effect on the evoked peak amplitude described for the AVREC waveform is reflected by the RMS amplitude in granular input layers. Here, ketamine anesthesia also led to an increase across all stimulation frequencies. Infragranular layers Vb and VIa show no difference of the evoked RMS (Figure 5, *top*). In contrast, the longer peak latencies of the AVREC waveform are reflected mainly in the temporal activity of supragranular layers. Here, peak latencies are significantly longer in the awake animals for all stimulation frequencies, except of the BF (Figure 5, *bottom*). Peak latencies of other sink components show only minor differences. In accordance with longer peak latencies for off-BF stimulation, the corresponding RMS amplitudes in supragranular layers are also significantly higher (Figure 5A, *top left*). This analysis of layer-specific sink activity reveals that the main drive for the time-locked columnar activity and higher peak amplitudes under ketamine is a strong excitation in cortical input layers III/IV. The compensation of the comparable overall columnar current flow in the awake condition is guaranteed by less time-locked, but stronger recruitment of upper layers for stimulation frequencies not representing the BF in the subsequent columnar processing. The increase of layer III/IV RMS activity appears to be mostly a modulation of gain paired with only a minor difference in tuning sharpness between the awake and ketamine group.

**Figure 5.**
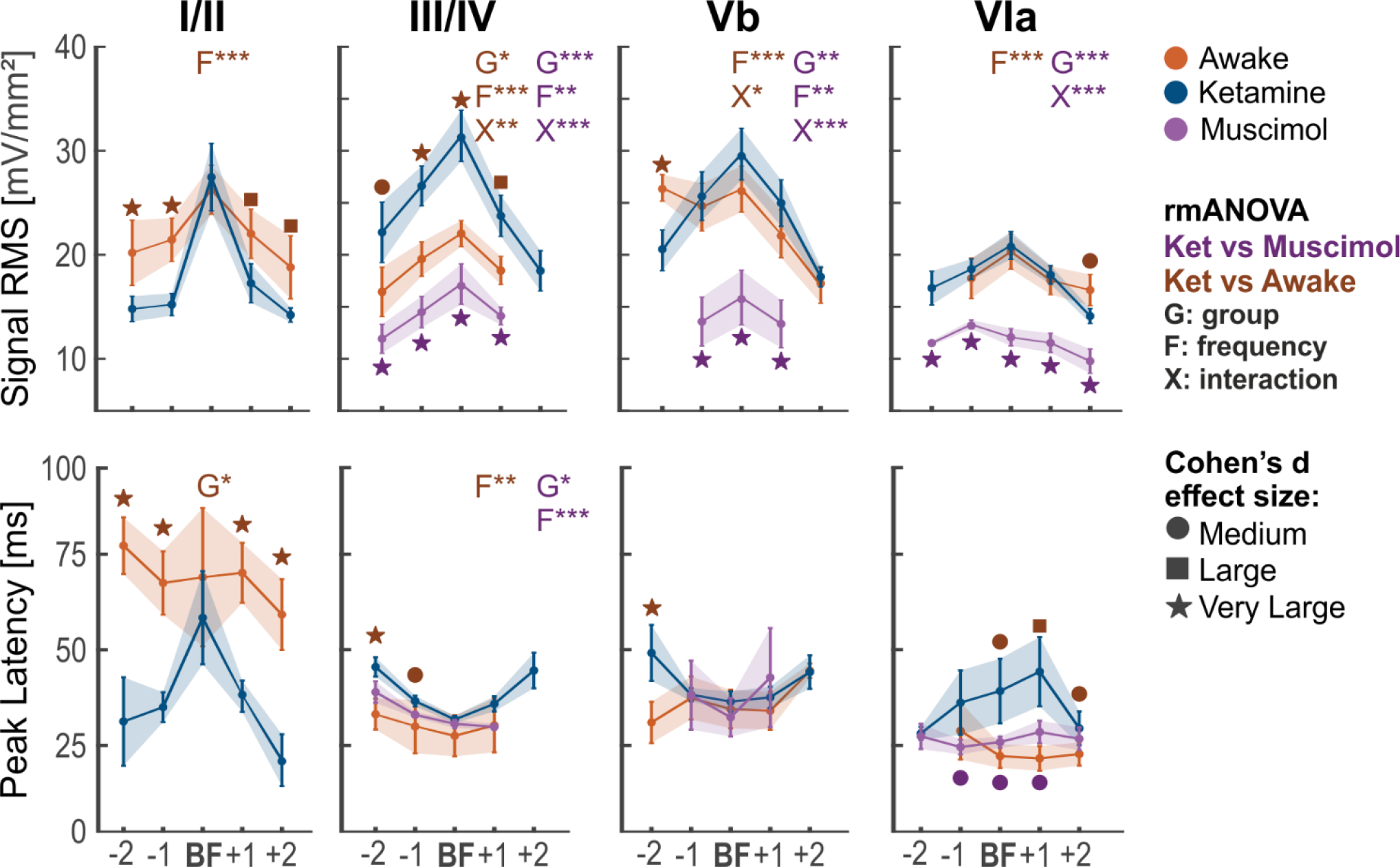
Sink tuning curves and peak latencies for early thalamic input layers. Semiautomatic detected sink activities within each layer. Tuning curves were sorted according to the layer-wise BF tuning for each layer for each group (orange: awake, n=9; blue: ketamine, n=11; purple: musicmol, n=11). Top, CSD signal associated RMS values. Bottom, Latency of peak response. Statistics are based on an rmANOVA, which shows group (G), frequency (F), or interaction (X) effects between both ketamine vs awake (orange) or ketamine vs muscimol (purple). *=p<0.05, **=p<0.01, ***=p<0.001. Cohen’s D indicates effect sizes with a circle for medium, a square for large, and a star for very large.

In order to differentiate effects of amplitude and phase-locking on the described gain effects implicated by our data, we used continuous wavelet analysis (Figure 6 and 7) at a laminar level to quantify the spectral representation between groups. In general, the spectral magnitude in granular layers III/IV show the highest difference between both groups with an increase in magnitude across all spectral bands in the ketamine group for BF stimulation (Figure 6A). This broadband significant increase between ketamine anesthetized and awake CSD scalograms was not present at BF-2, indicating more similar tuning curves for both groups at border frequencies (cf. Figure 5). The significant increase in magnitude for BF stimulation is revealed by a high Cohen’s D effect size and with a permutation clustermass test for all frequencies from theta (4-7 Hz) to high gamma (60-100 Hz). The nature of this broadband magnitude increase in cortical layers III/IV induced by ketamine is consistent with the immediate recruitment of synaptic activity indicating stimulus-locked gain increase. In other layers, magnitude effects are less pronounced. In Figure 6B, the supragranular I/II and infragranular VIa clustermass permutation tests reveal no areas of significant magnitude increase above chance. Layer Vb shows an increase across low and high beta and low gamma.

**Figure 6.**
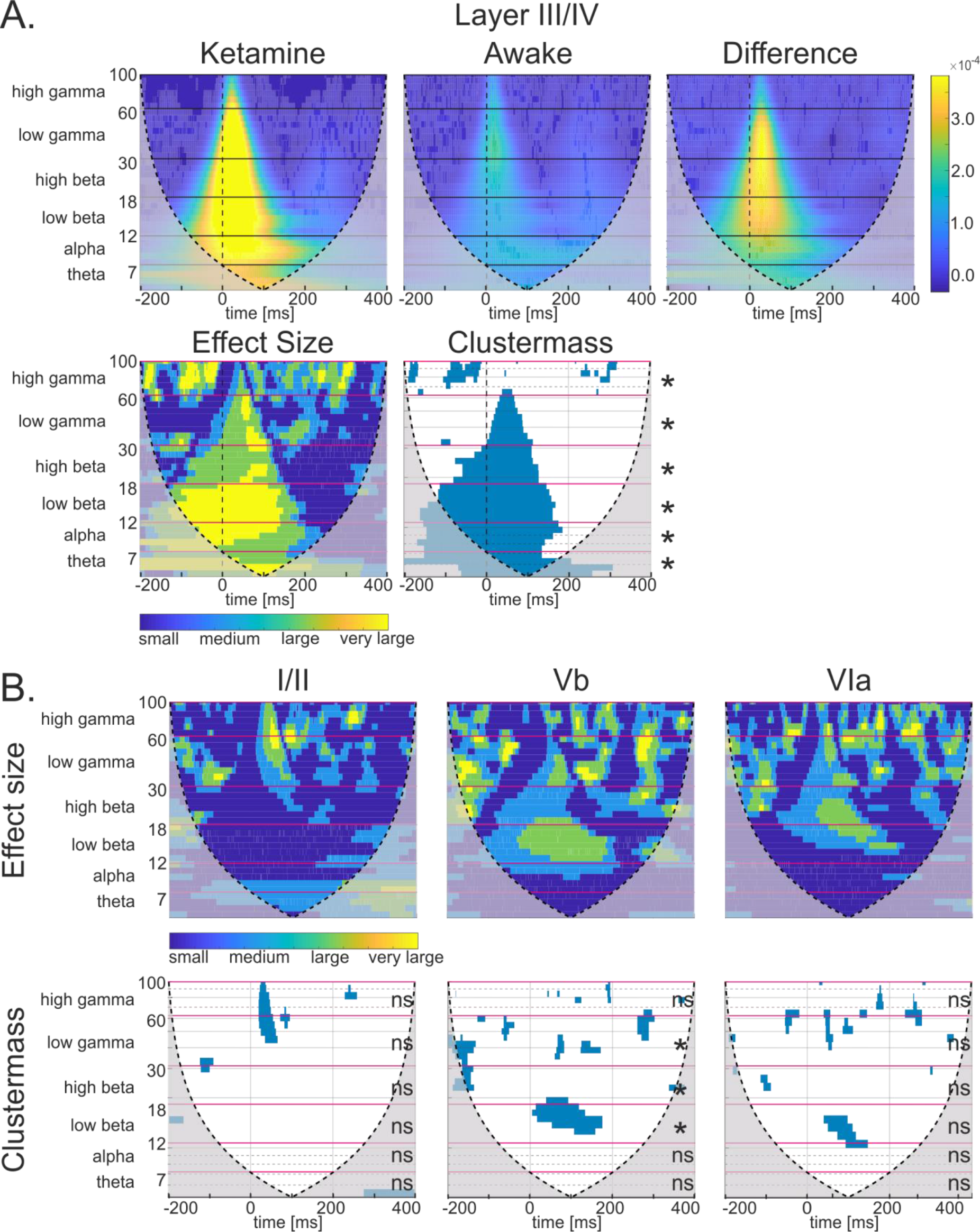
Magnitude scalograms of continuous wavelet transform and analysis from trial-averaged subject CSD profiles. ***A***, BF response of layer III/IV; scalograms of ketamine group (left, n=11), awake group (middle, n=9), and the absolute difference between both groups (right); Effect size (Cohen’s D) matrix showing small through very large effect size and clustermass matrix showing significance below p<0.05 (verified by two-sample student’s *t* test). Fuchsia lines indicate binning borders of frequencies for wavelet analysis and permutation test (left, *=p<0.05 or ns=not significant) of the clustermass. ***B***, Effect size (Cohen’s D) matrix, clustermass matrix, and permutation test results for layers I/II, Vb, and VIa at their respective BF. All graphs show the cone of influence overlaid as a dashed line and muted areas which extend outward. This shows where the wavelet transform was likely affected by boundary conditions.

**Figure 7.**
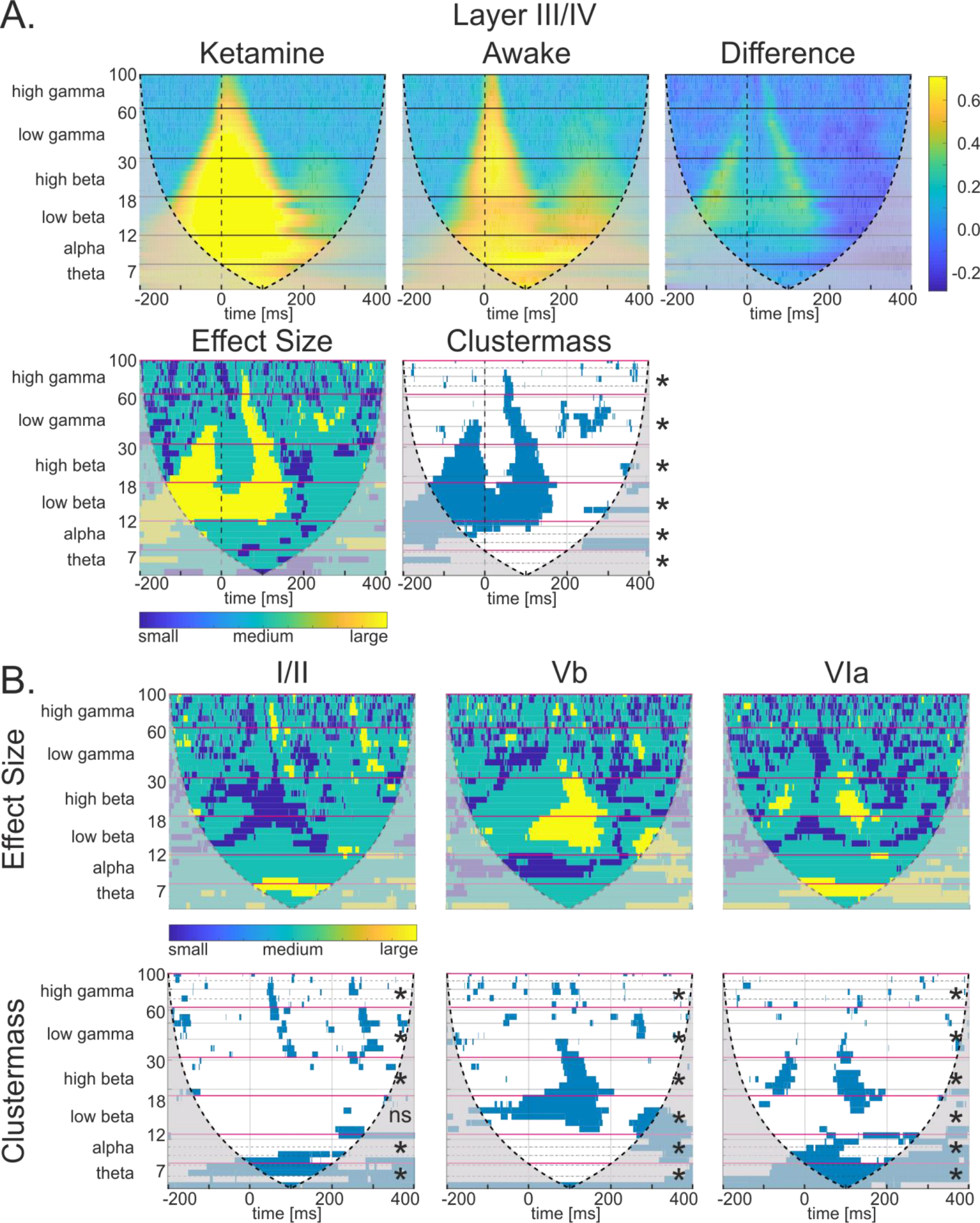
Phase coherence scalograms of continuous wavelet transform and analysis from single trial subject CSD profiles. ***A***, BF response of layer III/IV; scalograms of ketamine group (left, n=11), awake group (middle, n=9), and the absolute difference between both groups (right); Effect size matrix showing small through large effect size and clustermass matrix showing significance below p<0.05 (verified by two-sample student’s *t* test). Fuchsia lines indicate binning borders of frequencies for wavelet analysis and permutation test (left, *=p<0.05 or ns=not significant) of the clustermass. ***B,*** Effect size matrix, clustermass matrix, and permutation test results for layers I/II, Vb, and VIa at their respective BF. All graphs show the cone of influence overlaid as a dashed line and muted areas which extend outward. This shows where the wavelet transform was likely affected by boundary conditions.

Further, we analyzed single trial phase coherence for the corresponding individual layer-wise CSD traces (Figure 7). While ketamine increased the stimulus-induced magnitude most prominently at stimulus onset, its effects on phase coherence were mainly surrounding the time of stimulus-induced columnar response. A corresponding broadband increase in phase coherence across all frequency bands as well as a strong increase in the lower beta range, proceeded during stimulus presentation above chance according to permutation (Figure 7A). Beta phase coherence was also significantly increased above chance in thalamocortical input layer Vb. Effects were less prominent in cortical layers I/II and VIa (Figure 7B).

Finally, cortical layers III/IV reveal the highest level of a stimulus-locked broadband magnitude increase induced by ketamine, which we show is due to early-onset broadband increase in amplitude rather than stimulus-induced phase locking. However, ketamine also creates a longer stimulus-induced broadband phase coherence in granular layers. Our findings might be explained by an input-circuit specific dephasing of tone-evoked recurrent excitation and balanced inhibition to incoming external stimuli.

## Discussion

We investigated the effects of anesthetic doses of ketamine (15mg kg^−1^/h) on synaptic population dynamics in the auditory cortex of Mongolian gerbils using layer-specific CSD analysis. We intended to illuminate functional mechanisms under ketamine anesthesia which would benefit translational research on using it as a therapeutic agent. Our data revealed a gain increase in layer III/IV which was ascribed to an increase in tone-evoked amplitude across spectral frequencies in a continuous wavelet analysis (CWT) analysis rather than stimulus-response phase locking. Ketamine produced an area of higher phase coherence to incoming stimuli mainly in granular layers in comparison to a more variable signal-response in awake animals, but this difference was not significant during early thalamocortical input processing. We observed higher-stimulus-locked responses at more variable amplitudes under ketamine, specifically in granular thalamic input layers. Our findings therefore argue for an altered input processing due to an increase in recurrent excitation of thalamocortical inputs selectively in granular layers III/IV which may be attributed to a disinhibition of GABAergic interneurons, leading to a reduced coupling of excitatory and inhibitory input circuits under ketamine.

### Ketamine induces a higher time-locked stimulus response at more variable peak amplitudes

Ketamine reduces the variability of early thalamocortical input in granular layers, which accounts for the stimulus-locked response strength and, in turn, the reduced temporal variability of overall columnar activity. The grand average CSD profiles (Figure 1) show a visually distinguishable difference in the temporal flow of cortical processing between ketamine-treated and awake animals. CSD profiles after cortical silencing with topical application of muscimol demonstrate the early and short synaptic response of thalamic input in layers III/IV and Vb/VIa (Happel et al., 2010). The residual sinks after cortical silencing are shorter in duration than before muscimol application. Due to this, much of the following signal represented in the other two conditions can be considered cortical in nature rather than of thalamic origin. This reveals that cortical processing contributes to the peak and duration of early signal processing in the auditory cortex. This can be attributed to recurrent excitation of the stimulus response in cortical layers III/IV (Liu et al., 2007). Activity under ketamine, as well as ketamine and topical muscimol, is shown to be fairly stimulus-locked at the area of thalamic input while awake animals have a wider spread of signal response activity across time and space.

To validate that the effects we found across the cortex and in specific layers are significant above chance (Figure 2), a clustermass permutation test was performed between awake and ketamine anesthetized subjects. This revealed significance above chance between groups at a columnar level and most significantly in cortical layer III/IV across early and late time bins. We used analysis of the AVREC in order to determine differences of the overall, temporal columnar response profile (Figure 3). Single-trial data revealed a significantly higher peak amplitude and faster peak latency (Figure3B). Due to the difference in the temporal fine structure of the averaged AVREC traces between both groups (Figure 3A), we used the Brown-Forsythe test of variance at the single trial level to observe differences in the stimulus-induced peak responses. The variance of the peak latency of induced activity while awake was significantly higher than under ketamine anesthesia, whereas the opposite was found for the variance of peak amplitudes (Figure 4). This increased peak latency variability particularly indicates a different recruitment of cortical cell populations under ketamine, where probabilistic response dynamics are lost.

### Increased recurrent excitation in early granular activity under ketamine

Ketamine increases recurrent excitation in granular layers after thalamic input to the cortex. Layer-specific tuning curves (Figure 5) were calculated from a semi-automatic sink detection algorithm in order to further classify the differences found at the columnar level between groups. Significance was revealed in the RMS value around the stimulus response of early layer III/IV between awake and ketamine anesthetized gerbils. In contrast, the other layers receiving thalamic input, Vb and VIa, did not show any significant difference. We infer based on this result that the input from the thalamus is similar between awake and ketamine-anesthetized gerbils, which is in accordance with a recent study investigating whole-cell recordings with different sound levels and not finding differential thalamocortical input between conditions (Zhou et al., 2014). Therefore, the increased strength seen in granular input layers under ketamine anesthesia is likely cortically driven—possibly due to ketamine-induced inhibition of GABAergic populations in the granular recurrent feedback loop. Some single and multi-unit studies using ketamine anesthesia have also specifically implicated a lack of inhibitory modulations from supragranular populations to this recurrent excitation (Kato 2017; Wehr & Zador, 2003). This is in line with influences of ketamine on the excitatory and inhibitory balance via selectively reducing activity of PV-releasing GABAergic interneurons. Our data therefore supports the hypothesis of ketamine-driven cortical disinhibition (Miller et al., 2016) and is in support of previously revealed elevated levels of neural activity after ketamine due to increased cerebral blood flow (Långsjö et al., 2005).

### Rampant recurrent excitation of granular thalamic inputs mediates gain enhancement

To further explore the phenomenon of layer selective effects, we performed a CWT analysis separately for CSD traces from the supragranular I/II and thalamic input III/IV, Vb, and VIa layers. CWT analysis is ideally suited to analyze the amplitude and phase effects of the peak structure of evoked responses. Performing a clustermass permutation test to compare spectral representations in the awake and ketamine group, we demonstrated a significant magnitude broadband increase in response to pure-tone stimuli selectively in granular input layers under ketamine. Ketamine induced an increased broadband area of cross-trial phase coherence compared to awake cortices with the exception of the time range of early thalamocortical input processing. Therefore, the observed granular layer gain increase under ketamine is mainly due to amplitude effects and not to an increased stimulus-induced phase-locking (Figures 6 and 7). Together with the less variable tone-evoked peak latencies after ketamine (Figure 4), our analysis reflects a highly stimulus-locked recruitment of the synaptic circuits in layers III/IV dominating the columnar response to the BF stimulus. For stimulation frequencies at BF-2, effects were less prominent. This indicates that the input gain of the recurrent excitation in the granular layer, which is most effectively recruited by salient thalamic input, is the most likely network mechanism by which ketamine affects sensory processing in the cortex. Furthermore, the increased time-frequency area of increased phase coherence across trials that preceded the phase of dominant thalamocortical input processing may implicate a possible de-coupling effect of excitatory and inhibitory synaptic input due to a suppressed GABAergic inhibitory response. Thereby, cortical disinhibition may lead to a less effective balanced inhibition of thalamocortical inputs in A1 (cf. Wehr & Zador, 2003). This may further account for the variability of amplitude seen in the AVREC curves and the immediate, stimulus-locked recruitment of excitatory populations and therefore the lack of probabilistic peak response dynamics.

## Conclusion

Previous findings investigating ketamine *in vivo* have described differential effects based on brain region (Slovik et al., 2017; Widman & McMahon, 2018), method of investigation (Bojak et al., 2013; Hildebrandt et al., 2017; Schwertner et al., 2018), and emphasized the dose dependency of the observed effects (Ahnaou et al., 2017; Hertle et al., 2016; Macdonald et al., 1987). These diverse observations have reported frequency-specific increases or decreases also for the beta, theta and alpha band. However, the most robust finding throughout the literature is the increased gamma oscillation across regions and ketamine doses. We have now revealed in our study, that at least for cortical networks, this effect is most robustly explained by a rampant recurrent micro-excitation in granular input layers. The effect was most prominent during best-frequency stimulation and less specific for other cortical layers and stimulation frequencies. We therefore propose that effects on lower frequency bands, as also observed across layers and different stimulation frequencies in our study, strongly depend on the very nature of the cortical layer under investigation and type of stimulation. Future studies may therefore enunciate the specificity of higher gamma oscillations for the broadly reported psychological effects of ketamine, for instance for the development of new therapeutic agents with reduced side effects.

## Author contributions

Research was designed by KED and MFKH. MFKH supervised the project. Experiments have been performed by KED, MGKB, JM, XL, MMZ, FA, and SA. Experiments have been supervised by MFKH and MD. Data have been analyzed by KED, MGKB, AWC, and MMZ. Statistical analysis was done by KED. Figures were prepared by KED and AWC. KED wrote the initial paper draft. Manuscript was written by KED and MFKH with the assistance from MKB, MD and FWO. All authors reviewed the manuscript.

## Acknowledgements

We would like to thank Kathrin Ohl and Silvia Vieweg for their technical assistance. This project was founded by the Deutsche Forschungsgemeinschaft (DFG SFB 779) and the Leibniz Association by the LIN Postdoctoral Network (LPN) and the China Scholarship Council.

